# Nucleus accumbens acetylcholine receptors modulate the balance of flexible and inflexible cue-directed motivation

**DOI:** 10.1101/2022.12.22.521615

**Authors:** Erica S. Townsend, Kenneth A. Amaya, Elizabeth B. Smedley, Kyle S. Smith

## Abstract

Sign-tracking is a conditioned response where animals interact with reward-predictive cues and can be used as a means to approximate a cue’s motivational value. The nucleus accumbens core (NAc) has been highly implicated in mediating the sign-tracking response. Additionally, acetylcholine (ACh) transmission throughout the striatum broadly has been attributed to both incentive motivation and behavioral flexibility. Here, we show that sign-tracking responses are indeed flexible in the face of a contingency change in the form of an omission schedule, and that this flexibility is mediated by NAc ACh. Using behavioral and pharmacological methods, we show that blockade of NAc nicotinic receptors (nAChRs) augmented sign-tracking persistence, while blockade of muscarinic receptors (mAChRs) enhanced response flexibility following introduction of the omission schedule. Further, we detail how mAChR or nAChR antagonism impacted the microstructure of sign-tracking responses. These results indicate that NAc ACh receptors have opposing roles in the regulation of sign-tracking response flexibility without altering the motivational value of the cue.

## Introduction

Animals can attribute motivational value, or incentive salience, to conditioned stimuli (CS) that predict rewards in their environment [1]. This CS attraction can manifest as appetitive approach and engagement with the cue as if it were the reward itself, also known as sign-tracking [2–4]. Motivational attraction toward reward-predictive cues is evolutionarily adaptive; for example, using environmental cues to guide the pursuit of resources such as food and water. However, cue-reward relationships can change in realistic and dynamic environments which could promote maladaptive, or even compulsive, reward seeking if animals are inflexible in their pursuit of reward. Indeed, animals who tend to sign-track are notably vulnerable to making suboptimal choices and can exhibit addiction-like behaviors [5–7].

Despite this, sign-tracking is known to be adaptable when individuals are faced with changes in natural environments. Sign-tracking responses are robust, but animals can readily adapt their behavior to meet changes in outcome value [8,9] and outcome contingencies [10]. Notably, animals can adjust their responses to cues in changing cue-reward conditions but still maintain an attraction to them [10–13]. In a typical Pavlovian autoshaping procedure with a lever CS, sign-tracking behaviors often result in lever deflections. Upon introduction of a contingency change in which lever-cue deflection cancels reward delivery (“omission”), animals will gradually reduce their cue contact. However, this does not mean that the animal has ceased sign-tracking. Instead, animals continue engaging with the cue by flexibly updating their sign-tracking responses through distinct cue-directed behaviors that are less vigorous and thus do not result in deflections (e.g., orienting and sniffing) [10–13]. In sum, the cue’s incentive salience is preserved as evidenced by animals’ continued motivational persistence to the cue, yet behavioral flexibility is exhibited through their change in responding after omission learning. The omission schedule thus provides a unique window to study brain mechanisms for how reward cues compel an enduring attraction from animals, even when the animal must adapt to changing environments.

The nucleus accumbens core (NAc) has long been implicated in motivation and the sign-tracking response [14–19]. The region hosts a small population of tonically active, modulatory cholinergic interneurons (ChIs) that are the primary souce of acetylcholine (ACh) in the striatum [20]. As such, ACh receptors are densely expressed throughout the NAc and consist of two major subtypes: ionotropic nicotinic receptors (nAChRs; [21]) and G-protein coupled muscarinic receptors (mAChRs; [22]). nAChRs in the NAc are almost exclusively expressed on dopaminergic terminals from the ventral tegmental area (VTA; [23,24]), while mAChRs are primarily autoreceptors, located on ChIs themselves [25]. ACh has an important role in the modulation of dopamine (DA) signaling by acting as a filter; when ACh signaling is lower, more DA can be released from VTA terminals [23,25–29]. Thus, proper activity at ACh receptors is crucial for maintaining both tonic and phasic DA dynamics in the NAc, likely including cue-evoked DA release that is required for the maintenance and acquisition of the sign-tracking response [19].

Striatal ChIs and ACh transmission have been implicated in behavioral flexibility, motivationally driven responding, and context-related behavioral changes [26,30–37]. In the NAc specifically, blockade of nAChRs or mAChRs augments or reduces, respectively, cue-evoked behavior invigoration, motivation, and phasic DA in a Pavlovian-to-Instrumental transfer (PIT) task [35]. However, the role of ACh transmission in the flexibility of cue-directed motivation through sign-tracking is unknown. Based on previous work, It is thought that ACh may serve to modulate motivation itself [35]. However, sign-tracking, as a behavior, can provide us with a distinct perspective to disentangle motivation from action and delve into the influence of ACh on the flexibility of an animal’s motivational pursuit. To examine this, we introduced sign-tracking rats to a cue contingency change through an omission schedule. For the first five days of the omission schedule, the rats received intra-NAc infusions of mecamylamine or scopolamine, nAChR and mAChR antagonists, respectively. We found that rats infused with mecamylamine exhibited greater lever cue engagement and food-cup entries in response to the omission schedule, creating a new topography of their sign-tracking response that was distinct from that of control animals. These behavioral changes were exclusive to the omission task and were not seen in control groups where rats were infused with mecamylamine during sign-tracking overtraining. Conversely, scopolamine decreased the persistence of behaviors that led to lever deflections and altered the structure of behavior with an increase in behaviors that did not cancel reward delivery, thus rendering animals more flexible towards changes in cue contingency during omission. These results highlight an opposing role for NAc ACh receptor regulation in stable motivation, in which nAChRs encourage and mAChRs resist flexible responding during cue and associative structure changes.

## Methods

### Experiment 1: Nicotinic Receptor Blockade During Introduction of an Omission Schedule

#### Rationale

Experiment 1 was designed to test the role of nAChRs in the NAc on how an attraction to reward cues can be flexible yet persistent when vigorous cue interaction becomes disadvantageous.

#### Subjects

Experimentally naïve, PN 70-90 male and female Long Evans rats were obtained from Charles River (n = 18; Charles River, Indianapolis, IN). Rats were single housed, and on a 12 h light/dark cycle (lights on at 7:00 AM). Experiments were conducted during the light cycle. Rats were food restricted (7-15g of standard chow per day) to 85% of their free-feeding weight prior and throughout testing. During the post-operative period, food was available ad libitum. Water was available ad libitum throughout the duration of the experiments. All procedures were approved by the Dartmouth College Institutional Animal Care and Use Committee.

#### Surgical Procedures

Surgeries were conducted under aseptic conditions. Rats were anesthetized with isoflurane gas and placed in a stereotaxic apparatus (Stoelting, Kiel, WI). 22-gauge, stainless steel guide cannulas (P1 Technologies, Roanoke, VA) were implanted bilaterally, 1 mm above the intended NAc infusion site (AP +1.3 mm, ML ± 1.8, DV −6.2 relative to Bregma). These coordinates were decided upon based on a similar study assessing the role of NAc cholinergic transmission on motivation [35]. Following surgery, rats were given intraperitoneal (IP) injections of 3 mg/kg of ketoprofen, 3 mL of 0.9% sterile saline, and 0.02 mL of enrofloxacin antibiotics. Animals were allowed to recover for 5 days with ad libitum food and water. and given IP injections of sterile saline and enrofloxacin during the first 3 recovery days. Food restriction resumed a minimum of 5 days before behavioral procedures resumed.

#### Testing Apparatus

All behavioral training and testing were conducted in identical standard operant chambers (20 × 30.5 × 29 cm; Med Associates, St. Albans, VT) enclosed in sound- and light-attenuating cabinets (62 × 56 × 56 cm) equipped with an exhaust fan for airflow and background noise (~68 dB) and illuminated by a house light on the back wall. Chambers contained two retractable levers on either side of a recessed magazine in which food rewards were delivered. All lever depressions and magazine entries were recorded automatically using the MED-PC IV software (Med Associates, St. Albans, VT).

#### Sign-Tracking Training

Training began with a 30-minute magazine acclimation session in which pellets were delivered at a rate of approximately one pellet every 30 seconds. Rats then received 12 total days of Pavlovian sign-tracking (ST) training sessions. The first 10 sessions of Pavlovian training were given over 10 consecutive days, and following surgery and post-operative procedures, the rats received 2 more consecutive training sessions, acting as a reacquisition period. A given ST training session contained 25 CS+ trials in which a 10-second presentation of a retractable lever was followed by noncontingent 45 mg grain pellet (BioServ, Frenchtown, NJ) delivery through the magazine, and 25 CS-trials in which the 10-second presentation of the other retractable lever was followed by nothing. CS+ and CS-levers were counterbalanced across animals and trials were pseudorandomized in such a way that no more than two of the same type of trial was followed in sequence, and intertrial intervals had a length of approximately 2 min. These sessions spanned approximately 1 hour.

#### Omission Testing

After completing 12 days of training, the rats underwent 7 days of omission testing. Similar to the Pavlovian training schedule, these sessions contained 25, 10-second CS+ trials and 25, 10-second CS-trials. Under the omission condition, a deflection of any given CS+ lever during the 10-second presentation would result in no reward, or in other words, cancellation of reward delivery for that trial. CS+ trials in which the rats did not deflect the lever were rewarded.

#### Drug and Infusion Procedures

To habituate animals to infusion procedures, animals were handled by experimenters for one week before the start of all experimental procedures, in addition to the days leading up to infusion procedures post-surgery recovery. Prior to each of the first 5 omission testing sessions, rats were bilaterally infused with either mecamylamine (10 ug/side; Tocris Bioscience, Bristol, UK), a nonselective nicotinic receptor antagonist, or an equivalent volume of sterile artificial cerebrospinal fluid (ACSF; Tocris Bioscience). Rats were gently held while either the drug or ACSF were infused into the NAc at a volume of 0.5 uL over 1 min through an injector (P1 Technologies, Roanoke, VA) inserted into the guide cannula, protruding 1 mm ventral of the cannula tip. The injector rested in the cannula for 1 min after the infusion, and rats were kept in a holding chamber for 10 min before beginning the behavioral task.

#### Histological Procedures

Following testing in all experiments, rats were anesthetized with sodium pentobarbitol (100 mg/kg) and perfused intracardially with 0.9% saline, followed by 10% formalin. Brains were removed and stored in 20% sucrose for 24 hours, then sectioned at 60 uL. Sections were mounted on microscope slides and cover slipped with a DAPI-containing mounting medium (Vectashield; Vector Laboratories, Burlingame, CA, USA) to verify the placements of the cannulas. Cannula placement maps were created by manually transcribing the most ventral affected regions onto printed images, and then transcribed digitally via Adobe Illustrator (26.0.1 Adobe Creative Cloud).

#### Data Analysis

Lever deflections and pellet magazine entries were recorded through MedPC. Lever press rates are calculated in presses per minute (PPM). PPM is calculated by dividing the total presses for a lever by the total minutes of lever availability. All statistical tests were carried out using R (version 4.2.2) [38]. Individual mixed models were analyzed with package “lme4” from CRAN [39] to analyze the effects of dependent variable responding (e.g., lever PPM, mean magazine entries) by fixed effects of experimental group and session while accounting for random effects of differences in individual starting values and learning rates over sessions. Categorical variables with multiple levels (e.g., group and lever type) were dummy coded to make predetermined comparisons between levels (e.g., CS+ vs CS-, treatment vs control). These models were created for three major phases in the experiment: sign-tracking training (days 1-12), omission testing (days 13-19), and drug infusion (days 13-17). Reported statistics include parameter estimates (β values), 95% confidence intervals (CI), and *p*-values (R; “lmerTest”) [40]. Linear mixed models were used because they consider aspects of the data structure that repeated measures ANOVA cannot and allows for safer generalization to larger populations of animals. Plots were created using the “ggplot2” package for R [41], and formatted for publication in Adobe Illustrator.

Videos were hand-scored for 4 notable sessions: session 12 (last sign-tracking acquisition training session), 13 (first omission testing session), 17 (last infusion session), and 19 (last day of testing sessions). Within each session for a given animal, a sampling of 6, 10-second CS+ trials were scored, specifically the 1^st^, 5^th^, 10^th^, 15^th^, 20^th^, and 25^th^ trials. Behaviors that occurred on the odd seconds of the trial were recorded for a total of 30 behaviors scored per session, per animal. Behaviors were scored into one of 7 categories: lever bites, lever grabs (one paw on each side of the lever), lever contacts (one-pawed touch on any side of the lever), lever sniffs (snout close to or touching the lever), lever orients, magazine-directed behaviors (magazine entry or orienting towards the magazine), and other non-CS+ directed behaviors (orienting or approaching CS-lever wall, orienting away from CS+ lever, ignoring CS+ lever). Linear mixed models were analyzed using the “lme4” package from CRAN [39], similarly to lever deflection and magazine entry models. These models analyzed the effects of dependent variable responding (e.g., lever bites, lever grabs, and all other scored behavior categories) by fixed effects of experimental group and session while accounting for individual starting values. Reported statistics include parameter estimates (β values), 95% confidence intervals (CI), and *p*-values (R; “lmerTest”) [40]. Bar plots were created using the “ggplot2” package for R [41]. To explore data further and between groups in multiple experiments, correlation matrices for each group were created using the “cor” function in R, and correlation plots were created using the “corrplot” package for R [38]. Of note, “non-CS+ directed behaviors” were not included in correlation matrices as some groups exhibited zero of this type of response during sessions. All plots were formatted in Adobe Illustrator.

### Experiment 2: Muscarinic Receptor Blockade During Introduction of an Omission Schedule

#### Rationale

Experiment 2 was designed to test the role of mAChRs in the ability for sign-tracking microstructures to flexibly modify following a contingency change.

#### Subjects, behavioral training, surgical, and histological procedures

Subjects were 18 experimentally naïve male and female Long Evans rats obtained from the same vendor as Experiment 1 and maintained in the same conditions. The apparatus was the same as in Experiment 1. The rats underwent 1 day of magazine training, 12 days of sign-tracking training, and 7 days of omission testing. Surgical and histological procedures were as described in Experiment 1.

#### Omission Testing and Drug Infusion Procedures

After completing 12 days of training, the rats underwent 7 days of omission testing as described in Experiment 1. Prior to the first 5 omission testing sessions, rats were bilaterally infused with either scopolamine (10 ug/side; Tocris Bioscience, Bristol, UK), a nonselective muscarinic receptor antagonist, or an equivalent volume of sterile artificial cerebrospinal fluid (ACSF; Tocris Bioscience). All drug infusion and behavioral testing procedures were identical to those described in Experiment 1.

### Experiment 3: Nicotinic Receptor Blockade During Sign-Tracking Overtraining

#### Rationale

Mecamylamine affected the persistence and strategy of responding during omission in Experiment 1. It remained unclear whether it would affect the sign-tracking response without the introduction of the omission schedule. Experiment 3 addresses this by infusing mecamylamine during sign-tracking overtraining.

#### Subjects, behavioral training, surgical, and histological procedures

Subjects were 18 experimentally naïve male and female Long Evans rats obtained from the same vendor as Experiments 1 and 2 and maintained in the same conditions. The apparatus was the same as in Experiments 1 and 2. The rats underwent 1 day of magazine training and 12 days of sign-tracking training, Surgical and histological procedures were as described in Experiment 1.

#### Overtraining and Drug Infusion Procedures

After completing 12 days of sign-tracking training, the rats underwent 7 days of overtraining in which they continued the sign-tracking training program described in Experiments 1 and 2.

Infusion procedures proceeded as described in Experiments 1 and 2, thus, drug infusions were given for only the first 5 of 7 days of overtraining.

## Results

### Experiment 1: Nicotinic Receptor Blockade During Introduction of an Omission Schedule

To compare group press rates with respect to time during the three major phases of the experiment, linear mixed models used PPM as the dependent variable by fixed effects of session, group, and the interaction between session and group, with random intercepts for individual animal start points included (Figure 1C). The models indicated that all rats acquired the sign-tracking response at similar rates during the 12 sessions of training, showing no main effect of group (est: −0.20 PPM; CI: −2.79 — 2.38; p = 0.878), session (est: 1.41 PPM; CI: −0.05 — 2.87; p = 0.058), or session and group interaction (est: 1.25 PPM; CI: −0.21 — 2.71; p = 0.093). These data suggest that animals are statistically similar upon beginning pharmacological manipulations. A series of 7 omission test days followed, in which mecamylamine was infused for the first 5 sessions. During these five days, a significant main effect of group (est: 2.28 PPM; CI: 0.06 — 4.50; **p = 0.044**), indicating differences between mecamylamine and ACSF group PPM rates during the drug infusion period. There was also a significant main effect of session (est: −2.91 PPM; CI: −4.98 0.84; **p = 0.006**), but no significant interaction between session and group (est: −1.30 PPM; CI: −3.37 — 0.77; p = 0.217) was found. All animals decreased pressing over the 5 days, however the mecamylamine group had higher press rates in contrast to the ACSF group. Over the whole 7 sessions of omission testing, there was a main effect of session (est: −3.00 PPM; CI: −4.56 1.43; **p = <0.001**), indicating a significant decrease over the omission period, but no significant effects of group (est: 1.67 PPM; CI: −0.48 — 3.83; p = 0.128) or significant interactions between group and session (est: −1.43 PPM; CI: −3.00 — 0.13; p = 0.073) were found. This may suggest that drug effects are transitory, and do not impact behavior long term or when drugs are not infused.

**Figure 1:**
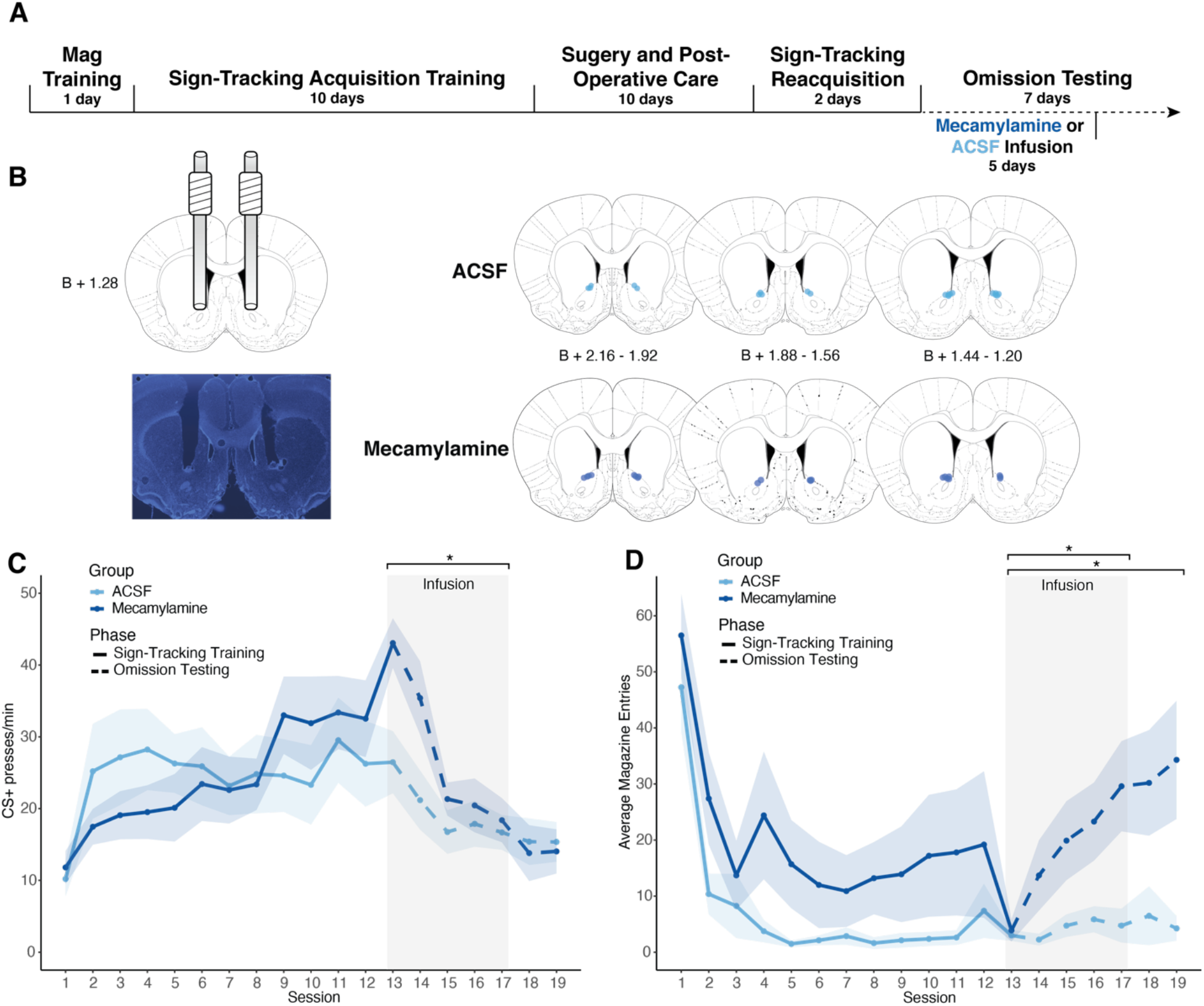
Experiment 1 results. (A) Timeline of behavioral training, surgical procedures, and infusions. (B) Cannula implant histology. All points mapped indicate the position of the bottom of the cannula, in the A/P coordinate closest to the center of the cannula implantation site. (C) Presses per minute (PPM) on the CS+ lever over the 12 sign-tracking training sessions (sessions 1-12, solid line) and 7 omission testing sessions (sessions 13-19, dotted line) for the mecamylamine (dark blue) and ACSF (light blue) groups. Shaded sessions indicate infusion sessions. Asterisks indicate significant (p < 0.05) linear mixed models over those sessions. (D) Average magazine entries during 10-second CS+ presentations over the 12 sign-tracking training sessions (sessions 1-12, solid line) and 7 omission testing sessions (sessions 13-19, dotted line) for the mecamylamine (dark blue) and ACSF (light blue) groups. Shaded sessions indicate infusion sessions. Asterisks indicate significant (p < 0.05) linear mixed models over those sessions. For all graphs, lines show mean and error shows ± SEM.

To compare mean group magazine entries with respect to time during the three major phases of the experiment, linear mixed models used the total magazine entries during only CS+ presentations as the dependent variable by fixed effects of session, group, and the interaction between session and group, with random intercepts for individual animal start points included (Figure 1D). During sign-tracking training, the model indicated a main effect of session (est: −6.91 entries; CI: −10.01 3.81; **p = <0.001**), but not group (est: 5.79 entries; CI: −1.99 — 13.57; p = 0.144) or group and session interactions (est: −0.60 entries; CI: −3.70 — 2.50; p = 0.705). All animals decreased magazine entries during cue presentations across training, and both groups did so similarly. During the five sessions of drug infusion, main effects of session (est: 4.84 entries; CI: 2.18 — 7.51; **p = 0.001**), group (est: 6.98 entries; CI: 1.14 — 12.82; **p = 0.020**), and a significant interaction between group and session (est: 3.83 entries; CI: 1.17 — 6.50; **p = 0.005**) were found. This indicates that magazine entries of animals in the mecamylamine group increased at a much steeper rate than animals in the ACSF group. Main effects of session (est: 5.24 entries; CI: 2.13 — 8.35; **p = 0.001**), group (est: 8.82 entries; CI: 1.81 — 15.84; **p = 0.014**), and a significant interaction between group and session (est: 4.36 entries; CI: 4.36; CI: 1.25 — 7.47; **p = 0.006**) were also found throughout the 7 omission testing sessions. This demonstrates that experimental group animals increased their magazine entries during the CS+ significantly throughout the whole omission testing phase.

### Experiment 2: Muscarinic Receptor Blockade During Introduction of an Omission Schedule

Press rates were compared with respect to session during the three major phases of the experiment using linear mixed models. These models used lever presses per minute as the dependent variable by fixed effects of session, group, and the interaction between session and group (Figure 2C). Random intercepts of individual animal start points were included. During the first 12 days of sign-tracking acquisition training, the models indicated a significant effect of session (est: 1.80 PPM; CI: 0.32 – 3.29; **p = 0.018**), but no significant effect of group (est: −1.08 PPM; CI: −4.58 – 2.41; p = 0.542) or interaction between group and session (est: −0.55 PPM; CI: −2.03 – 0.94; p = 0.470). This indicated that both groups acquired the sign-tracking response similarly prior to any manipulations. During scopolamine infusion, the models showed a significant effect of session (est: −2.38 PPM; CI: −4.31 – −0.45; **p = 0.016**) and group (est: −3.07 PPM; CI: −6.00 – −0.14; **p = 0.040**). No significant interaction between session and group were found (est: 1.15 PPM; CI: −0.77 – 3.08; p = 0.239). All animals therefore decreased lever deflections over these sessions, and scopolamine group animals were lower overall in contrast to animals in the ACSF group. Over the whole 7 days of omission testing, a significant effect of session was seen (est: −2.11 PPM; CI: −3.62 – −0.60; **p = 0.006**), but no significant effect of group (est: −2.42 PPM; CI: −5.10 – 0.27; p = 0.078) or interaction between session and group were seen (est: 1.43 PPM; CI: −0.08 – 2.94; p = 0.063). This may suggest that drug effects are transitory and exclusive to infusion sessions, and do not impact behavior long term.

**Figure 2:**
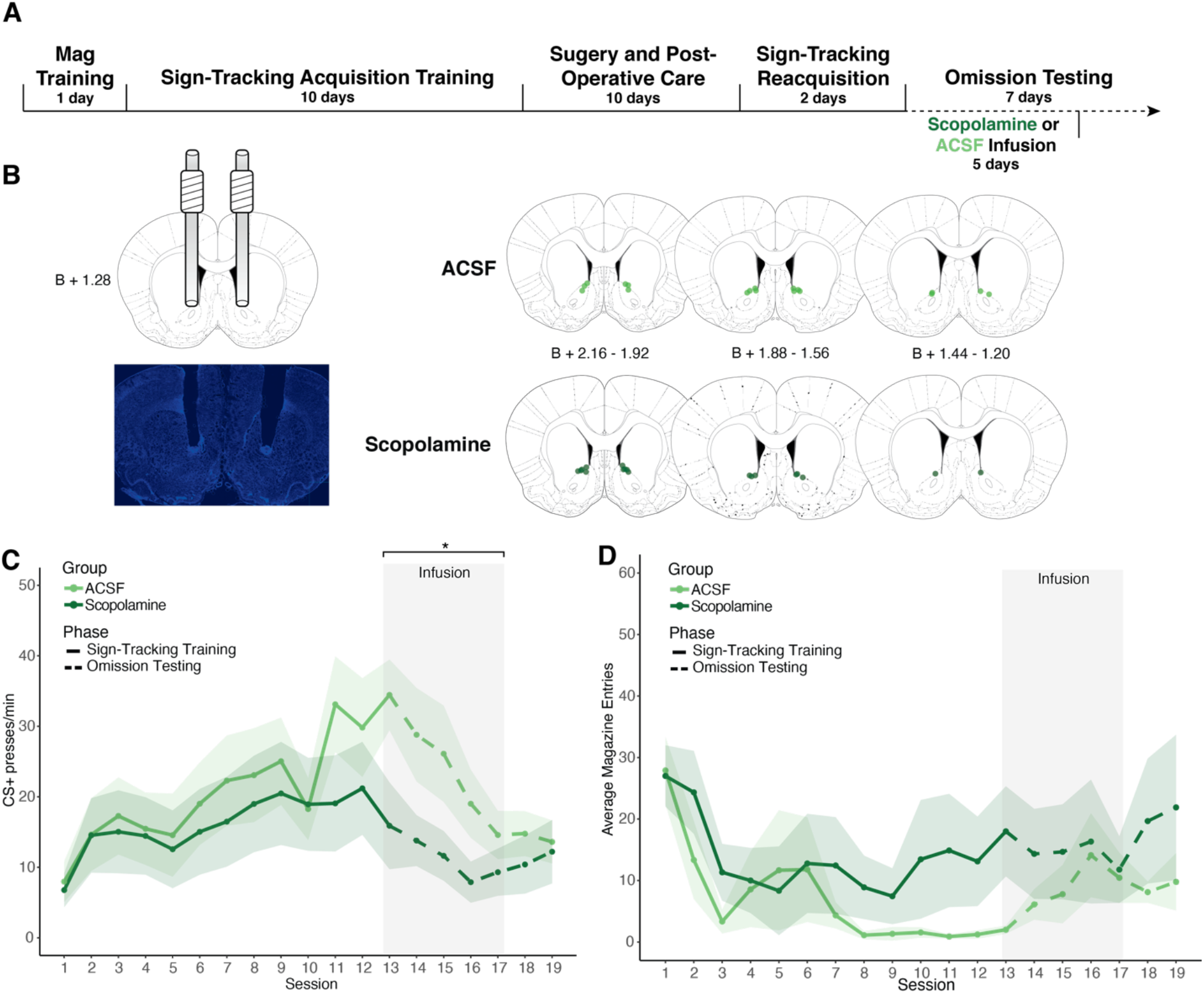
Experiment 2 results. (A) Timeline of behavioral training, surgical procedures, and infusions. (B) Cannula implant histology. All points mapped indicate the position of the bottom of the cannula, in the A/P coordinate closest to the center of the cannula implantation site. (C) Presses per minute (PPM) on the CS+ lever over the 12 sign-tracking training sessions (sessions 1-12, solid line) and 7 omission testing sessions (sessions 13-19, dotted line) for the scopolamine (dark green) and ACSF (light green) groups. Shaded sessions indicate infusion sessions. Asterisks indicate significant (p < 0.05) linear mixed models over those sessions. (D) Average magazine entries during 10-second CS+ presentations over the 12 sign-tracking training sessions (sessions 1-12, solid line) and 7 omission testing sessions (sessions 13-19, dotted line) for the scopolamine (dark green) and ACSF (light green) groups. Shaded sessions indicate infusion sessions. Asterisks indicate significant (p < 0.05) linear mixed models over those sessions. For all graphs, lines show mean and error shows ± SEM.

To compare magazine entries between groups with respect to time during the three major phases of the experiment, linear mixed models used the total magazine entries during only CS+ presentations as the dependent variable by fixed effects of session, group, and the interaction between session and group, with random intercepts for individual animal start points included (Figure 2D). During the first 12 days of sign-tracking acquisition training, the models indicated no significant effects of session (est: −0.54 entries; CI: −5.69 – 4.61; p = 0.837), group (est: −0.87 entries; CI: −7.70 – 5.96, p = 0.803), or significant interactions between session and group (est: −1.63 entries; CI: −6.78 – 3.52; p = 0.534). No significant effects of session (est: 1.88 entries; CI: −0.27 – 4.03; p = 0.085), group (est: −3.97 entries; CI: −16.32 – 8.37; p = 0.526), or an interaction between the two (est: −0.93 entries; CI: −3.08 – 1.22; p = 0.393) were found during scopolamine infusion. During the whole 7 days of omission testing, there were also no effects of session (est: 1.88 entries; CI: −0.27 – 4.03; p = 0.085), group (est: −3.97 entries; CI: −16.32 – 8.37; p = 0.526), or an interaction between session and group (est: −0.93 entries; CI: −3.08 – 1.22; p = 0.393). Overall, no changes in magazine entries during cue presentations were prevalent.

### Experiment 3: Nicotinic Receptor Blockade During Sign-Tracking Overtraining

To compare group press rates with respect to session during the three major phases of the experiment, linear mixed models used PPM as the dependent variable by fixed effects of session, group, and the interaction between session and group, with random intercepts for individual animal start points included (Figure 3C). The models indicated that all rats acquired the sign-tracking response at similar rates during the 12 sessions of training, showing a significant main effect of session (est: 4.46 PPM; CI: 2.89 — 6.03; **p = <0.001**), and no main effects of group (est: 0.89 PPM; CI: −2.09 — 3.87; p = 0.557) nor a significant session and group interaction (est: 0.99 PPM; CI: −0.59 — 2.71; p = 0.218). A series of 7 overtraining sessions followed in which there was no task change (i.e., no omission). Mecamylamine was infused for the first 5 of these overtraining sessions. During these five sessions, the model indicated no significant effects of group (est: 3.92; CI: −0.65 — 8.49; p = 0.092), session (est: −0.98 PPM; CI: −4.01 — 2.04; p = 0.522), nor interactions between session and group (est: −0.24 PPM; CI: −3.26 — 2.79; p = 0.876). Over the total 7 sessions of overtraining, no effects of session (est: −1.28 PPM; CI: −3.78 — 1.21; p = 0.313), group (est: 3.07 PPM; CI: −1.56 — 7.70; p = 0.192) nor significant interactions between group and session (est: −1.22 PPM; CI: −3.71 — 1.28; p = 0.337) were found. This indicates no significant effects of mecamylamine infusion on sign-tracking responses, and thus motivation generally, during overtraining when no task change was present.

**Figure 3:**
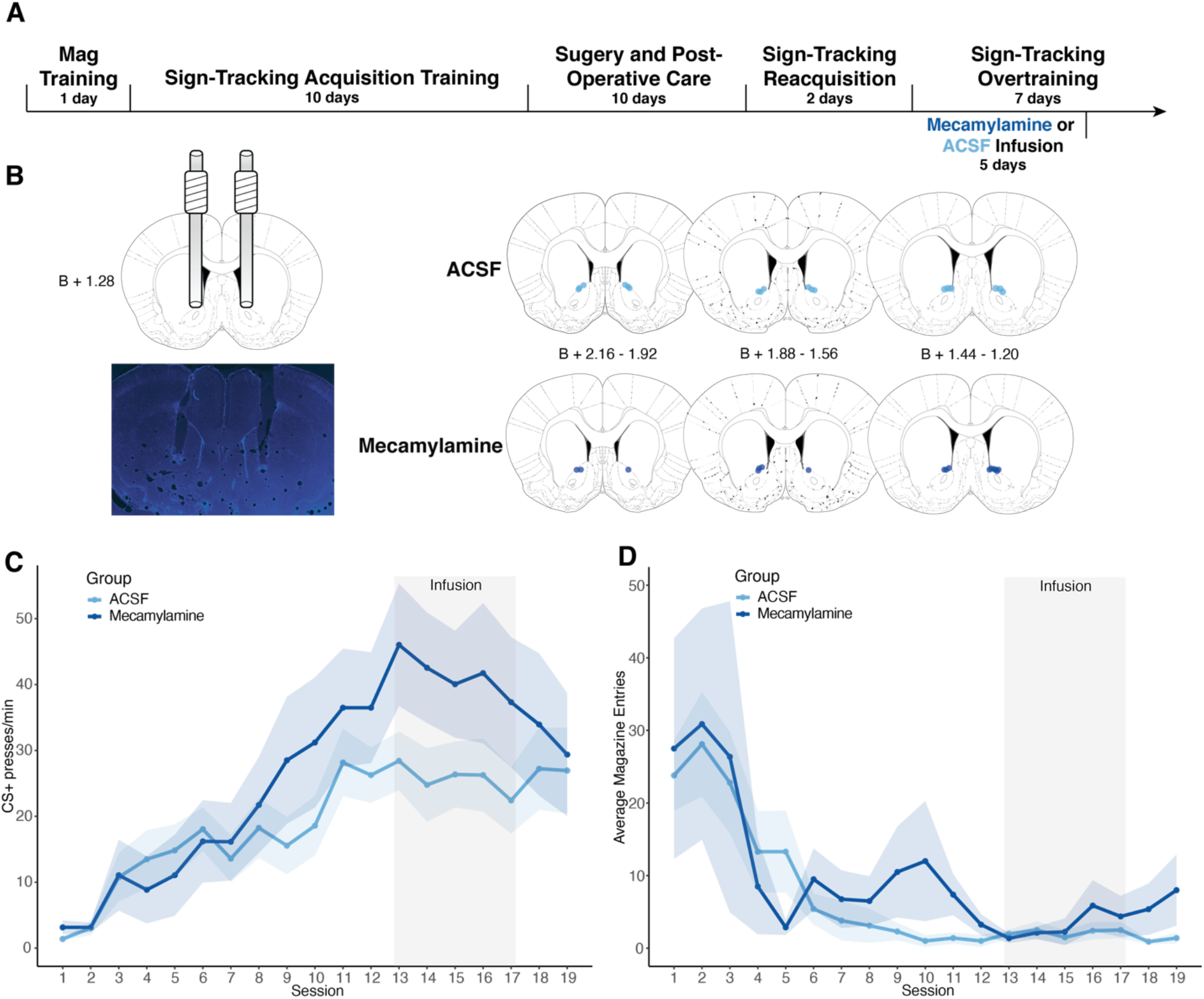
Experiment 3 results. (A) Timeline of behavioral training, surgical procedures, and infusions. (B) Cannula implant histology. All points mapped indicate the position of the bottom of the cannula, in the A/P coordinate closest to the center of the cannula implantation site. (C) Presses per minute (PPM) on the CS+ lever over the 12 sign-tracking training sessions (sessions 1-12) and 7 sign-tracking overtraining sessions (sessions 13-19) for the mecamylamine (dark blue) and ACSF (light blue) groups. Shaded sessions indicate infusion sessions. Asterisks indicate significant (p < 0.05) linear mixed models over those sessions. (D) Average magazine entries during 10-second CS+ presentations over the 12 sign-tracking training sessions (sessions 1-12) and 7 sign-tracking overtraining sessions (sessions 13-19) for the mecamylamine (dark blue) and ACSF (light blue) groups. Shaded sessions indicate infusion sessions. Asterisks indicate significant (p < 0.05) linear mixed models over those sessions. For all graphs, lines show mean and error shows ± SEM.

To compare mean group magazine entries with respect to time during the three major phases of the experiment, linear mixed models used the total magazine entries during only CS+ presentations as the dependent variable by fixed effects of session, group, and the interaction between session and group, with random intercepts for individual animal start points included (Figure 3D). During sign-tracking training, the model indicated a main effect of session (est: −7.84 entries; CI: −12.91 2.78; **p = 0.003**), but not group (est: 1.23 entries; CI: −3.18 — 5.65; p = 0.582) or group and session interactions (est: 0.67 entries; CI: −4.40 — 5.74; p = 0.795). During the five sessions of drug infusion, no significant main effects of session (est: 0.76 entries; CI: −0.33 — 1.85; p = 0.171), group (est: 0.51 entries; CI: −1.18 — 2.20; p = 0.551), nor a significant interaction between group and session (est: 0.63 entries; CI: −0.46 — 1.72; p = 0.254) were found. No main effects of session (est: 0.88 entries; CI: −0.44 — 2.20; p = 0.190), group (est: 1.16 entries; CI: −0.96 — 3.27; p = 0.282), nor a significant interaction between group and session (est: 1.17 entries; CI: −0.15 — 2.48; p = 0.083) were found throughout the 7 overtraining sessions. This demonstrates that there were no effects of mecamylamine on magazine entries during the CS+ presentations.

### Experiments 1 and 2 Response Microstructure Analysis

To compare the changes in the microstructure of sign-tracking responses between control and experimental groups from the last day of sign-tracking training through the end of omission testing (specifically, days 12, 13, 17, and 19 of the task), linear mixed models were employed using the mean of each given behavioral response type (lever bite, lever grab, lever contact, lever sniff, lever orient, magazine-directed, and non-CS+ directed; Figure 4) as the dependent variable by fixed effects of session, group and the interaction between session and group, with random intercepts for individual animal start points included. In Experiment 1, significant effects between groups were seen only in the lever contact and the magazine-directed behavior models (Figure 4A). The lever contact model showed a significant effect of group (est: −0.97 lever contacts; CI: −1.81 0.13; **p = 0.024**), and the magazine-directed behavior model showed a significant effect of group (est: 1.43 magazine-directed behaviors; CI: 0.36 – 2.49; **p = 0.010**) and a significant interaction between group and session (est: 0.82 magazine-directed behaviors; CI: 0.16 – 1.48; **p = 0.015**). In Experiment 2, significant effects of group were seen in models predicting lever bites (est: −1.35 lever bites; CI: −2.64 – −0.05; **p = 0.042**), lever grabs (est: −1.64 lever grabs; CI: −2.86 – −0.42; **p = 0.009**), lever orients (est: 0.51 lever orients; CI: 0.05 − 0.97; **p = 0.029**), and lever sniffs (est: 1.15 lever sniffs; CI: 0.08 – 2.22; **p = 0.035**; Figure 4B). Linear mixed models for other behaviors in Experiments 1 and 2 did not show significant effects of group or significant interactions (see Figure 4 and Supplementary Table 1).

**Figure 4:**
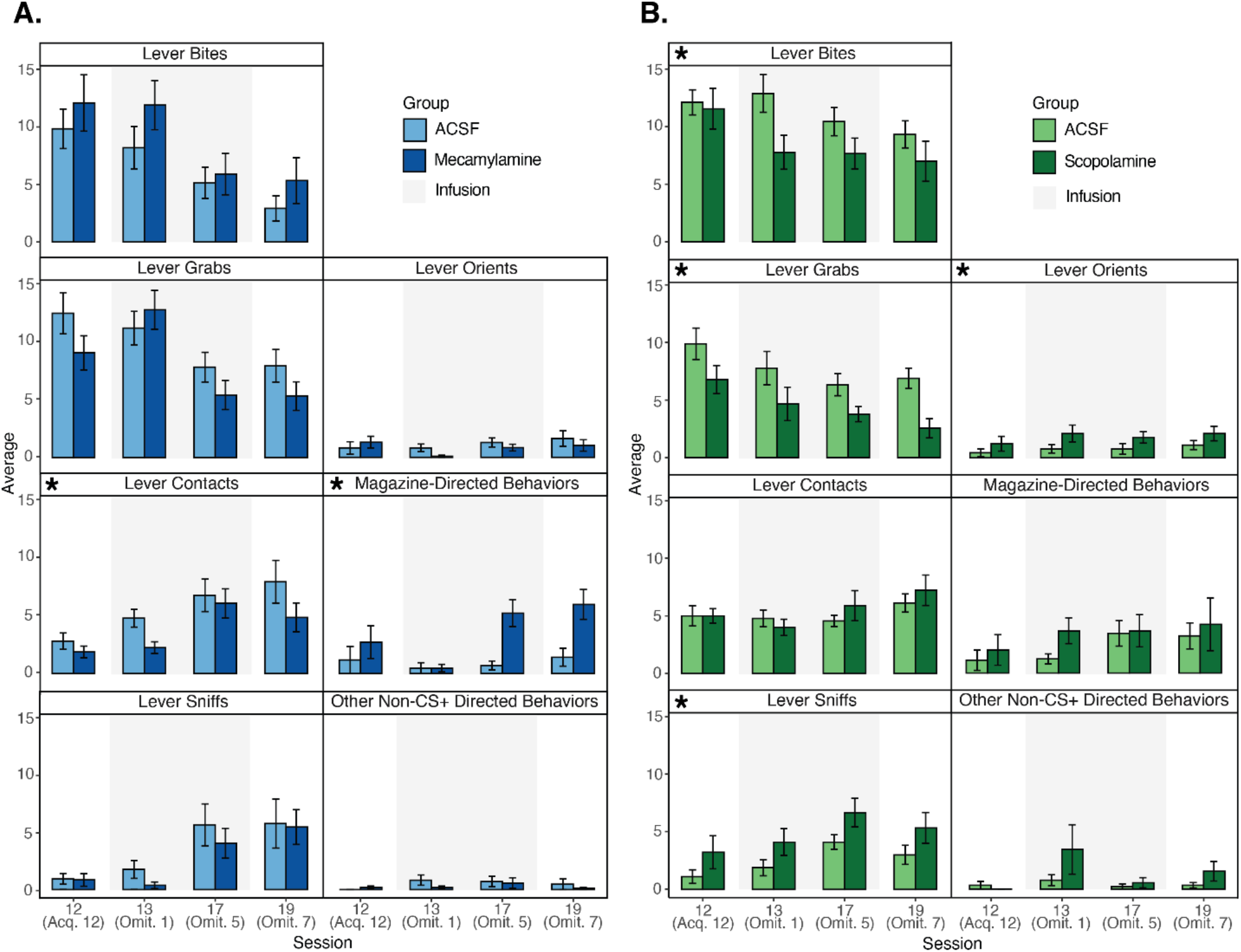
Microstructure of behavior. (A) Average scored behaviors exhibited during 10-second CS+ presentations during Experiment 1 in the mecamylamine (dark blue) and ACSF (light blue) groups. Infusion sessions are indicated by gray shading. (B) Average scored behaviors exhibited during 10-second CS+ presentations during Experiment 2 in the scopolamine (dark green) and ACSF (light green) groups. Infusion sessions are indicated by gray shading. For all graphs, bars show the mean and error bars show ± SEM, and asterisks represent significant linear mixed model group and/or interaction effects.

To analyze and compare the patterns of animals in each experiment, correlation matrices were created, and subsequently correlated against each other, for each experimental group in both Experiments 1 and 2 (Figure 5). The highest correlation, and thus the highest similarity between groups, were shown between Experiment 1 and Experiment 2 ACSF control groups (r = 0.6220). Common clusters highly positively correlated between these ACSF groups occurred in lever bites, and magazine directed behaviors (Figure 5). These behaviors appeared to be relatively stable through the 4 sampled omission sessions due to the consistent correlations across sessions. Of note, lever grabs in the Experiment 2 ACSF group were a much more prominent cluster than in the Experiment 1 ACSF group. Notable inversely correlated clusters in the Experiment 1 ACSF group, between lever sniffs and lever bites, and lever bites and lever orients, also occurred in the Experiment 2 ACSF group, although these were not as consistent across the 4 sessions as in Experiment 1 controls. This indicates that, typically, when lever bites were less frequent, lever sniffs and orients were more frequent.

**Figure 5:**
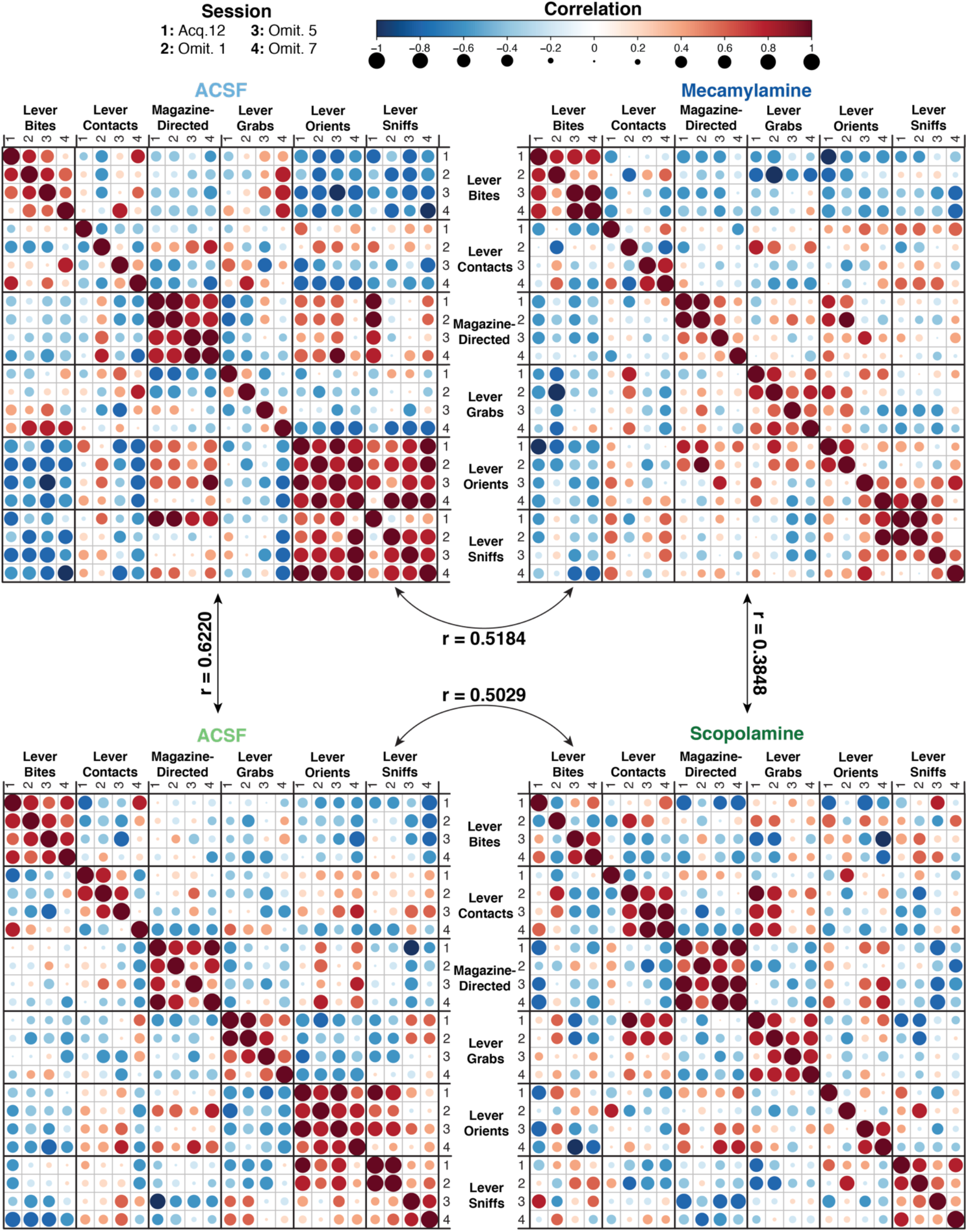
Patterns and correlation of behaviors in ACSF (top left) and mecamylamine (top right) groups in Experiment 1, and ACSF (bottom left) and scopolamine (bottom right) groups in Experiment 2. Correlations are represented by color and dot size. Correlations between groups are indicated by arrows between the matrices. Session number coding is as follows: 1 – Sign-Tracking Acquisition Session 12; 2 – Omission Session 1 (Session 13); 3 – Omission Session 5 (Session 17); 4 – Omission Session 7 (Session 19).

The correlation between ACSF and mecamylamine groups in Experiment 1 (r = 0.5184), indicated less similarity between these two groups than between the two ACSF groups. Similar positively correlated clusters between these groups included lever bites, as well as magazine-directed behaviors. However, lever bites tended to be more consistently stable across sessions in the mecamylamine group and magazine directed behaviors tended to be less stable through the 4 sampled sessions in relation to controls, consistent with magazine entry data. The large positively correlated cluster between lever orients and lever sniffs in the ACSF group appeared to disintegrate in the mecamylamine group, with much weaker correlations throughout most sessions. Additionally, the negatively correlated cluster apparent in the ACSF group, between lever orients and lever sniffs, still existed but were not as clearly consistent and disintegrated in the mecamylamine group. Lower contact responses, such as lever sniffs and lever orients, did not characterize the responding of the mecamylamine group as well as the ACSF group. The few strongly positive clusters in the mecamylamine group existed in higher contact behaviors, such as lever bites and lever grabs, consistent with higher press rates. This demonstrated clear differences of the sign-tracking microstructures between experimental groups, particularly as the tendencies of mecamylamine animals were to engage in less low-contact behaviors (such as lever orienting and lever sniffing).

The correlation between ACSF and scopolamine groups in Experiment 2 (r = 0.5029) was lower than that of the two control groups, which indicated higher levels of dissimilarity between groups in Experiment 2. Clusters in the correlation matrix were similar, however the consistency over sessions was much greater in controls than in the scopolamine group, particularly lever bites and lever contacts. Additionally, in the ACSF group lever sniffs appeared as a part of a larger positive cluster along with lever orients. This cluster disintegrated in the scopolamine group, although not fully, as lever sniffs still appeared as a strongly positive cluster. The negatively correlated clusters in lever orients and lever sniffs that were seen in controls were not apparent in scopolamine, as many of these correlations became more positive. This indicated that during some sessions, these orient and sniff responses tended to be co-occurring. These findings in total demonstrated a more diverse array and variability of behaviors in scopolamine animals. Notably, they described deterioration in higher lever contact behaviors (such as lever biting), enhancing performance in omission testing.

The lowest correlation that was found was between the Experiment 1 and Experiment 2 drug groups (r= 0.3848), which indicated the lowest similarity between groups. Primary dissimilarities between these drug groups occurred within lever bites, magazine-directed behaviors, and lever sniffs and orients. Lever bites were less stable and magazine-directed behaviors were more stable across sessions in the scopolamine group in comparison to the mecamylamine group. Lever grabs were quite similar in correlation strength and in the stability over sessions between both groups. Both groups had disintegration with the notable lever orient and sniff cluster that was evident in both ACSF groups. However, the strong correlations in the experimental groups that remained within these behaviors were expressed differently across sessions. The inversely correlated cluster that was evident in lever orients and sniffs in the mecamylamine group (in addition to both ACSF groups), was not prominent in the scopolamine group, further indicating fundamental differences in behavioral response topography of these experimental groups. These results indicated diverging behavioral response structures between mecamylamine and scopolamine groups, just as these two experimental groups had opposing effects on lever deflection rate in comparison to ACSF.

## Discussion

Reward-predictive cues have a powerful ability to motivate behavior towards the cues themselves, termed sign-tracking behavior [1,4]. As the behavior develops, sign-tracking responses can be sensitive to the cue-reward relationship [1,3,9,10,42,43]. Yet, once sign-tracking is established, cue attraction can remain powerful and enduring in the face of changing cue-reward relationships. This is seen when sign-trackers are faced with an omission schedule as they initially continue to engage with the lever cue in ways that produce deflections, even when doing so cancels reward delivery. Ultimately, animals learn to interact with the lever in a way that allows rewards to be delivered by restructuring their cue-directed response to avoid lever deflections that would omit reward delivery. Within the brain, the NAc ACh system shows promise as a potential mechanism of not necessarily motivation itself, but maintenance of motivation when cue-reward relationships change. Appropriate responding requires a carefully modulated balance of both flexibility and persistence. Here, we have evaluated NAc ACh in the context of sign-tracking behavior during an omission schedule and found that activity at the two major subtypes of ACh receptors have significant and opposing functions in how animals restructure their sign-tracking when navigating an omission procedure.

Through these experiments, we tested the hypothesis that blocking NAc ACh activity at nAChRs or mAChRs would specifically affect sign-tracking flexibility (i.e., how animals restructure their interaction with reward cues during omission learning). Our results show that these two receptors have opposing influences on this type of motivational flexibility. The blockade of nAChRs augmented, while blockade of mAChRs reduced the persistence of lever deflections during omission. These response alterations were further characterized as alterations to flexible motivated responding, as changes to responses were not only seen both in the form of lever presses, but additionally within magazine entries and diverging differences in response structures in which each drug group was characterized by unique patterns of behaviors. Blockade of nAChRs resulted in augmented magazine entries and magazine-directed behaviors during the cue presentations, however changes in these behaviors were not apparent when mAChRs were blocked. This finding was unexpected, but it shows that animals with nAChR blockade changed their response strategy not only in cue-directed behavior but also in magazine-directed behavior as though exploring the magazine for potential rewards. Additionally, the microstructures of behavior in mecamylamine and scopolamine groups showed the lowest correlation to one another. This relatively low similarity between groups in the makeup of their behavioral responding underscores the idea that nAChR and mAChR receptors act in opposition. Importantly, nAChR blockade only augmented sign-tracking behaviors under an omission schedule and did not alter motivation broadly as seen in sign-tracking overtraining. This suggests that ACh activity, specifically at nAChRs, may support a flexibility or response updating mechanism rather than simple modulation of motivated responding to cues.

Based on this set of results, we suggest that ACh transmission can serve as a balancing mechanism for how flexible animals are in their attraction to reward cues, with the nAChRs biasing animals towards flexibility and the mAChRs biasing animals towards inflexibility.

Endogenous ACh release would affect this balance by acting on both receptors, but perhaps at different time courses or intra-striatal locations. In this line of reasoning, our results suggest to us that there are orthogonal brain mechanisms for cue-driven motivation and for how malleable that motivation is in the way it is expressed behaviorally. ACh may have a special role in the latter process, regulating how physical responses play out when one is attracted to reward-related stimuli. An implication of this idea is that excessive or otherwise rigid motivational pursuit of goals could come about either through heightened motivational states or else stable motivational states that lose amenability to change.

These results are consistent with prior work implicating NAc ACh receptor activity in opposing roles in cue-motivation in a Pavlovian-to-Instrumental Transfer task (PIT). In a study by Collins and colleagues, a learned Pavlovian light cue and instrumental lever cue are presented as a compound cue. An opposing role of nAChRs and mAChRs were found, by which compound cue lever presses were increased or decreased, respectively, when these receptors were blocked, indicating differential modulation of motivation [35]. The “transfer” tested in PIT could be viewed as a type of flexibility on its own – this compound cue is new to the animal, and they must integrate known information into a new cue context. In PIT, it may be difficult to disentangle whether animals are less motivated, or if they are integrating information into their actions differently when ACh transmission changes. Sign-tracking affords us a different perspective of fundamental motivation itself. Thus, our results as well as those obtained in PIT may be a result of a potential cue-information integration and motivational flexibility mechanism of NAc ACh. This is not to say that ACh never contributes to the basic aspects of motivated behavior and incentive salience, as striatal cholinergic interneurons have been shown to respond or pause firing during both cues and rewards [30,31,44–47]. Further, striatal ACh manipulations can alter motivated responding generally [33,36,37,48]. However, we raise the possibility that ACh may play a more preferential role in fine-tuning *how* motivated behaviors occur once they are established, more so than *whether* they will occur or not. This role in flexibility is similar to that seen in the striatum broadly [32,34,49–53], and could be a key to understanding how ACh can manage behavioral adjustments through response modulation.

ACh receptor activity could alter the flexibility of cue-motivated behaviors, including sign-tracking, through a few potential mechanisms. First, the effects of mecamylamine could have been the result of action at nAChRs on presynaptic dopaminergic terminals. Activity at these receptors is known to regulate behaviorally relevant dopaminergic activity and has been dubbed as a “low-pass filter” of high-frequency dopaminergic stimulations [29]. These receptors have also been shown to regulate cue-evoked phasic DA in response to salient events such as reward-predictive cues [54]. The location of these receptors places them in a prime spot for direct modulation of known incentive salience and reward prediction error signaling, potential neural mechanisms underlying this type of responding. It is also known that cholinergic interneurons pause their tonic firing during salient events, such as during rewards [30,31]. This pausing may allow for their tight regulation of DA to lift momentarily to allow for these types of signals [26].

NAc ACh and its known regulation of phasic DA release, situates it in a central position to alter DA transmission to support flexibility in dynamic environments, changing structures, and new contexts. This mechanism both maintains and regulates motivation, but also creates the fine balance in which optimum reward and persistence are mediated as well. ACh receptor activity could be a novel neural regulatory mechanism over cue responding in changing conditions when flexibility is necessary. Persistent maladaptive motivation responses and cue reactivity, such as those that occur in substance use disorders and behavioral addictions, could be regulated by such a mechanism. Cue exposure therapies have promising results in treating those with substance use disorders [55–57], but they often fail to produce substantial and lasting changes to real world cues in different contexts [58–61]. Thus, the findings that blockade of nAChRs augmenting persistence of sign-tracking under omission, while blockade of mAChRs reduce persistence, may be therapeutically relevant. These results may be implicated in treatment of substance use and motivation disorders through furthering our understanding of the neural mechanisms underlying cue learning and responding that exist in natural and dynamic environments.

## Author Contributions

EST, KAA, and EBS, designed the studies. EST collected and analyzed the data, created figures, and wrote the draft manuscript. KAA and EBS contributed to data analysis and draft manuscript revision. KSS provided funds and supervision, was involved in overall design of the project, and the writing of the manuscript.

