## Supplemental Tables for "Nucleus accumbens acetylcholine receptors modulate the balance of flexible and inflexible cue-directed motivation"

**Supplementary Materials:**

Supplementary Table 1:

| Lever Bites | | | |
|---|---|---|---|
| **Predictor** | **Estimate** | **CI** | **P-Value** |
| Group | 1.22 | -1.05 – 3.48 | 0.288 |
| Session | -3.00 | -3.95 – -2.06 | **< 0.001** |
| Group × Session | -0.15 | -1.09 – 0.80 | 0.756 |
| **Lever Grabs** | | | |
| **Predictor** | **Estimate** | **CI** | **P-Value** |
| Group | -0.92 | -2.34 – 0.51 | 0.203 |
| Session | -2.15 | -3.18 – -1.11 | **< 0.001** |
| Group × Session | -0.09 | -3.18 – -1.11 | 0.859 |
| **Lever Contacts** | | | |
| **Predictor** | **Estimate** | **CI** | **P-Value** |
| Group | -0.97 | -1.81 – -0.13 | **0.024** |
| Session | 1.82 | 1.01 – 2.63 | **< 0.001** |
| Group × Session | -0.28 | -1.09 – 0.53 | 0.496 |
| **Lever Sniffs** | | | |
| **Predictor** | **Estimate** | **CI** | **P-Value** |
| Group | -0.45 | -1.70 – 0.80 | 0.477 |
| Session | 2.14 | 1.38 – 2.90 | **< 0.001** |
| Group × Session | -0.05 | -0.81 – 0.71 | 0.894 |
| **Lever Orients** | | | |
| **Predictor** | **Estimate** | **CI** | **P-Value** |
| Group | -0.17 | -0.66 – 0.32 | 0.486 |
| Session | 0.17 | -0.08 – 0.42 | 0.177 |
| Group × Session | -0.18 | -0.43 – 0.07 | 0.151 |
| **Magazine-Directed Behaviors** | | | |
| **Predictor** | **Estimate** | **CI** | **P-Value** |
| Group | 1.43 | 0.36 – 2.49 | **0.010** |
| Session | 0.93 | 0.28 – 1.59 | **0.006** |
| Group × Session | 0.82 | 0.16 – 1.48 | **0.015** |
| **Non-CS+ Directed Behaviors** | | | |
| **Predictor** | **Estimate** | **CI** | **P-Value** |
| Group | -0.13 | -0.37 – 0.12 | 0.302 |
| Session | 0.08 | -0.15 – 0.32 | 0.480 |
| Group × Session | -0.07 | -0.31 – 0.16 | 0.541 |

**Table 1:** Linear mixed model estimates, confidence intervals (CI), and p-values for group, session, and group and session interaction predictors for the 7 behaviors scored in Experiment 1. Significant values are indicated in bold.

Supplementary Table 2:

| Lever Bites | | | |
|---|---|---|---|
| **Predictor** | **Estimate** | **CI** | **P-Value** |
| Group | -1.35 | -2.64 – -0.05 | **0.042** |
| Session | -1.38 | -2.24 – -0.52 | **0.002** |
| Group × Session | -0.17 | -1.03 – 0.69 | 0.697 |
| **Lever Grabs** | | | |
| **Predictor** | **Estimate** | **CI** | **P-Value** |
| Group | -1.64 | -2.86 – -0.42 | **0.009** |
| Session | -1.35 | -1.89 – -0.81 | **< 0.001** |
| Group × Session | -0.18 | -0.72 – 0.37 | 0.522 |
| **Lever Contacts** | | | |
| **Predictor** | **Estimate** | **CI** | **P-Value** |
| Group | 0.21 | -0.60 – 1.01 | 0.608 |
| Session | 0.66 | 0.12 – 1.20 | **0.018** |
| Group × Session | 0.31 | -0.23 – 0.85 | 0.262 |
| **Lever Sniffs** | | | |
| **Predictor** | **Estimate** | **CI** | **P-Value** |
| Group | 1.15 | 0.08 – 2.22 | **0.035** |
| Session | 0.94 | 0.39 – 1.50 | **0.001** |
| Group × Session | 0.06 | -0.50 – 0.61 | 0.840 |
| **Lever Orients** | | | |
| **Predictor** | **Estimate** | **CI** | **P-Value** |
| Group | 0.51 | 0.05 – 0.97 | **0.029** |
| Session | 0.24 | -0.07 – 0.55 | 0.121 |
| Group × Session | 0.02 | -0.29 – 0.33 | 0.904 |
| **Magazine-Directed Behaviors** | | | |
| **Predictor** | **Estimate** | **CI** | **P-Value** |
| Group | 0.57 | -0.94 – 2.08 | 0.454 |
| Session | 0.86 | 0.34 – 1.38 | **0.002** |
| Group × Session | -0.11 | -0.63 – 0.41 | 0.684 |
| **Non-CS+ Directed Behaviors** | | | |
| **Predictor** | **Estimate** | **CI** | **P-Value** |
| Group | 0.49 | -0.21 – 1.18 | 0.165 |
| Session | 0.07 | -0.52 – 0.66 | 0.816 |
| Group × Session | 0.13 | -0.46 – 0.72 | 0.658 |

**Table 2:** Linear mixed model estimates, confidence intervals (CI), and p-values for group, session, and group and session interaction predictors for the 7 behaviors scored in Experiment 2. Significant values are indicated in bold.
